# Cryo-EM structure of the rhodopsin-Gαi-βγ complex reveals binding of the rhodopsin C-terminal tail to the Gβ subunit

**DOI:** 10.1101/547919

**Authors:** Ching-Ju Tsai, Jacopo Marino, Ricardo J. Adaixo, Filip Pamula, Jonas Mühle, Shoji Maeda, Tilman Flock, Nicholas M.I. Taylor, Inayatulla Mohammed, Hugues Matile, Roger J. P. Dawson, Xavier Deupi, Henning Stahlberg, Gebhard F. X. Schertler

## Abstract

G protein-coupled receptors (GPCRs) are the largest class of integral membrane proteins and represent key targets for pharmacological research. GPCRs modulate cell physiology by engaging and activating a diversity of intracellular transducers, prominently heterotrimeric G proteins, but also G protein-receptor kinases (GRKs) and arrestins. The recent surge in the number of structures of GPCR-G protein complexes has expanded our understanding of G protein recognition and GPCR-mediated signal transduction. However, many aspects of these mechanisms, including the existence of transient interactions with transducers, have remained elusive.

Here, we present the cryo-EM structure of the light-sensitive GPCR rhodopsin in complex with heterotrimeric Gi. In contrast to all reported structures, our density map reveals the receptor C-terminal tail bound to the Gβ subunit of the G protein heterotrimer. This observation provides a structural foundation for the role of the C-terminal tail in GPCR signaling, and of Gβ as scaffold for recruiting Gα subunits and GRKs. By comparing all available complex structures, we found a small set of common anchoring points that are G protein-subtype specific. Taken together, our structure and analysis provide new structural basis for the molecular events of the GPCR signaling pathway.

## Introduction

G protein-coupled receptors (GPCRs) are the most diverse class of integral membrane proteins with almost 800 members in humans. GPCRs are activated by a great diversity of extracellular stimuli including photons, neurotransmitters, ions, proteins, and hormones (Glukhova et al., 2018). Upon activation, GPCRs couple to intracellular transducers, including four subtypes of G proteins (Gαs, Gαi/o, Gαq/11, Gα12/13) (Milligan & Kostenis, 2006), seven subtypes of GPCR kinases (GRKs) (Gurevich, Tesmer, Mushegian, & Gurevich, 2012), and four subtypes of arrestins (Smith & Rajagopal, 2016) (**Figure 1A**), among many other partners (Magalhaes, Dunn, & Ferguson, 2012). While most GPCRs are promiscuous and can couple to more than one G protein subtype (Flock et al., 2017), the molecular determinants of G protein recognition are not yet fully understood. Understanding the molecular basis for G protein coupling and selectivity could lead to the design of drugs that promote specific signaling pathways and avoid unwanted side effects (Hauser, Attwood, Rask-Andersen, Schiöth, & Gloriam, 2017).

**Figure 1.**
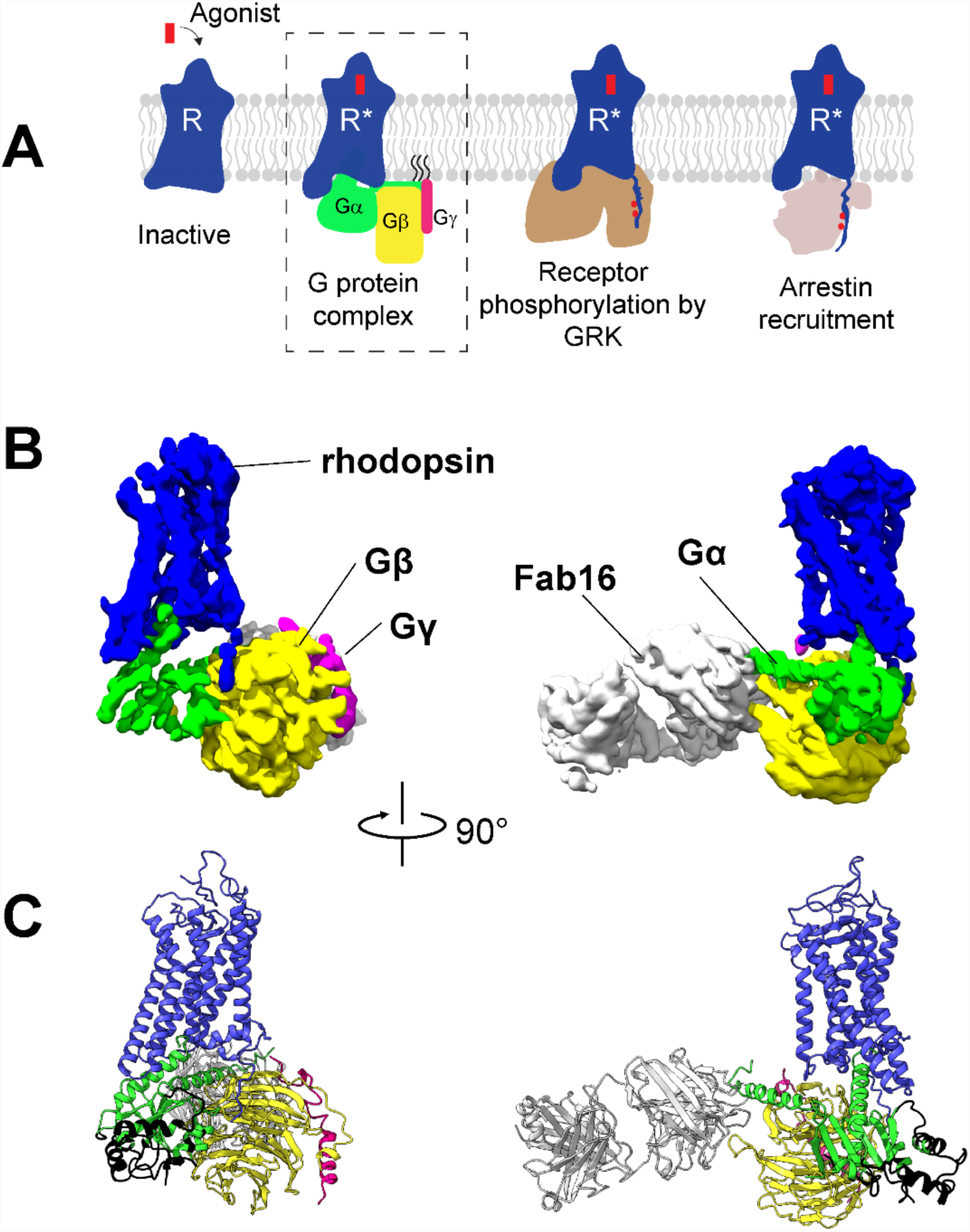
Cryo-EM structure of the rhodopsin-Gi-Fab16 complex. **(A)** GPCR signaling complexes. **(B)** EM density map of the complex (rhodopsin – blue, Gαi – green, Gβ – yellow, Gγ – magenta, Fab16 – white). **(C)** Atomic model of the complex (same color code as B). The black region of the Ras domain of Gα has no corresponding density in the EM map.

The recent surge in the number of structures of GPCR-G protein complexes has greatly expanded our understanding of G protein recognition and GPCR-mediated signal transduction. Out of the 13 structures of GPCR-G protein complexes available, six contain a Gi/o subtype: μ-opioid receptor bound to Gi (Koehl et al., 2018), adenosine A_1_ receptor bound to Gi (Draper-Joyce et al., 2018), cannabinoid receptor 1 bound to Gi (Krishna Kumar et al., 2019), human rhodopsin bound to Gi (Kang et al., 2018), 5HT_1B_ receptor bound to Go (García-Nafría, Nehmé, Edwards, & Tate, 2018), and bovine rhodopsin bound to an engineered Go (mini-Go) (Tsai et al., 2018). However, the preparation of GPCR-G protein complexes for structural biology still remains challenging (Munk et al., 2019). Nanobodies (Rasmussen et al., 2011), Fab fragments (Kang et al., 2018; Koehl et al., 2018), and mini-G proteins (García-Nafría, Nehmé, et al., 2018; Tsai et al., 2018) have been very important tools to overcome the inherent instability and flexibility of these complexes and obtain near atomic-resolution structures. Importantly, these structures represent snapshots of a particular state of the complex in the signaling cascade, and therefore additional structural data is required to improve our understanding of this process (Capper & Wacker, 2018).

The photoreceptor rhodopsin is one of the best-characterized model systems for studying GPCRs, providing invaluable information on how receptor activation is translated into G protein and arrestin binding and activation (K. P. Hofmann et al., 2009; L. Hofmann & Palczewski, 2015; Scheerer & Sommer, 2017). Rhodopsin has been shown to interact with the Gβγ subunit of the G protein heterotrimer to assist binding and activation of the Gα subunit (Herrmann et al., 2004, 2006). After dissociation of the Gαβγ heterotrimer, the Gβ subunit recruits GRKs to the membrane, resulting in the phosphorylation of the receptor C-terminal tail (C-tail) (Claing, Laporte, Caron, & Lefkowitz, 2002; Pao & Benovic, 2002; Pitcher, Freedman, & Lefkowitz, 1998) and binding of arrestin (Goodman Jr et al., 1996). Despite this biochemical evidence, a direct interaction between rhodopsin and Gβγ could not be observed in the existing complex (Kang et al., 2018).

Here, we present the cryo-EM structure of bovine rhodopsin in complex with a heterotrimeric Gi. Overall, our structure agrees well with a currently published structures (Kang et al., 2018; Tsai et al., 2018). Remarkably, the EM density map provides a first structural evidence for the interaction between the C-tail of the receptor and the Gβ subunit. The density map also shows that intracellular loops (ICL) 2 and 3 of rhodopsin are at contact distance to Gα. This prompted us to perform a comparison of all available structures of GPCR-G protein complexes to generate a comprehensive contact map of this region. We then extended this analysis to the binding interface formed by the C-terminal helix α5 of Gα and found that only a few G protein subtype-specific residues consistently bind to the receptors. These contacts are ubiquitous anchoring points that may be also involved in the selective engagement and activation of G proteins.

## Results

To obtain a rhodopsin-Gi complex suitable for structural studies, we expressed in HEK cells the constitutively active mutant of bovine rhodopsin N2C/M257Y/D282C (Deupi et al., 2012) which binds to the Gi protein heterotrimer (Maeda et al., 2014). The construct of the human Gαi1 subunit was expressed in *E. coli* (Sun et al., 2015), while the Gβ1γ1 subunit was isolated from bovine retinae and thus contains native post-translational modification important for transducin function (Matsuda & Fukada, 2000). We reconstituted the rhodopsin-Gαi1β1γ1 complex in the detergent lauryl maltose neopentyl glycol (LMNG) (Maeda et al., 2014) in presence of Fab16 (Maeda et al., 2018) **(Suppl. Figure 1)**, as cryo-EM screening revealed that the complex without Fab could not be refined to high resolution (**Suppl. Figure 2, Suppl. Table 1**). During unsupervised 3D classification of the complex bound to Fab16 (**Suppl. Figure 3**), we observed that the density corresponding to Gα is heterogenous **(Suppl. Figure 4)**, particularly at flexible regions such as the α-helical (AH) domain. The AH domain was then excluded by using a soft mask during refinement, resulting in a map with a nominal global resolution of 4.38 Å **(Suppl. Figure 3D, E)**. The EM map was used to build a model of the complex (**Suppl. Figure 5)**.

### Architecture of the rhodopsin-G protein complex

The structure of rhodopsin-Gi-Fab16 reveals the features observed in previously reported GPCR-G protein complexes **(Figure 1B, C; Suppl. Figure 6A)**. In particular, our cryo-EM structure is in excellent agreement with the crystal structure of the same rhodopsin mutant bound to a mini-Go protein (Tsai et al., 2018), with a nearly identical orientation of the C-terminal α5 helix **(Suppl. Figure 6B)**, which contacts transmembrane helices (TM) 2, 3, 5-7 and the TM7/H8 turn of the receptor **(Suppl. Figure 5B, and Suppl. Table 2)**.

### Interaction between the C-terminal tail of rhodopsin and Gβ

The EM map reveals a density on the Gβ subunit as continuation of H8 of the receptor **(Figure 2A)**, which corresponds to the C-tail of rhodopsin. This feature does not exist in any other GPCR-G protein complex, including the human rhodopsin-Gi complex bound to Fab_G50 (Kang et al., 2018). We modeled half of the C-tail of rhodopsin (12 out of 25 residues; 324-335) into this density as a continuous stretched peptide with residues G324, P327 and G329 serving as flexible hinges (**Figure 2B**). This allows us to compare the structure of this region in three signaling states of the receptor: inactive (Okada et al., 2004), G protein-bound (this structure), and arrestin-bound (Zhou et al., 2017) (**Figure 2B**). In the inactive state, the C-tail folds around the cytoplasmic side of rhodopsin, although this is likely due to crystal packing as this region is intrinsically disordered in the absence of a binding partner (Jaakola, Prilusky, Sussman, & Goldman, 2005; Venkatakrishnan et al., 2014). In our G protein complex, the C-tail stretches over the cleft between Gα and Gβ, interacting with both subunits (**Figure 2C**). In the rhodopsin-arrestin complex part of the proximal segment of the C-terminus (residues 325-329) is not resolved, but the distal part could be modeled up to S343, eight residues longer than in our G protein complex and including two phosphorylated sites. In the presence of arrestin, the C-tail stretches further, with residues K339-T342 forming a short β-strand antiparallel to the N-terminal β-strand of arrestin (**Figure 2B**). Thus, it appears that distinct portions of the C-tail are responsible for contacting different intracellular partners. The central part of the C-tail (residues 330-335) can bind to Gα and Gβ, while the distal half of the C-tail (residues 336-343), which contains five (out of six) phosphorylation sites (Azevedo et al., 2015), binds to arrestin (**Figure 2C**).

**Figure 2.**
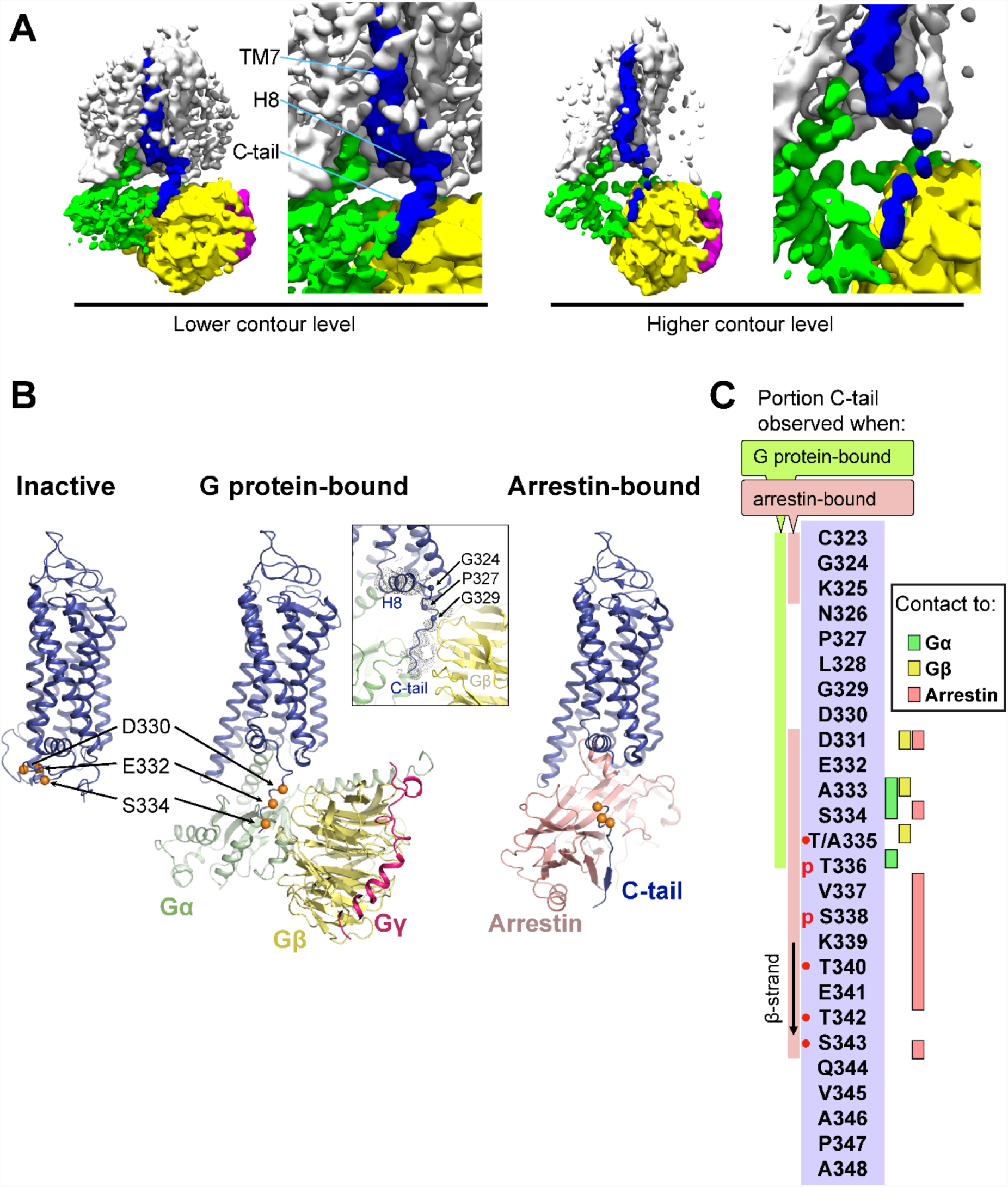
The C-terminal tail of rhodopsin. **(A)** The EM map is contoured at two different levels to show the continuity of the density. TM7, H8 and the C-tail of the receptor are colored in blue, Gα in green, Gβ in yellow, and Gγ in magenta. **(B)** Conformational change of the C-tail between three different signaling states of rhodopsin: Inactive state (left, PDB id: 1U19), G protein-bound (center, this work), and arrestin-bound (right, PDB id: 5W0P, chain A). The Cα atoms of residues Asp330, Glu332 and Ser334 are shown as orange spheres to help tracking the structural changes in the C-tail. All structures are aligned to rhodopsin. **(C)** Schematic representation of the rhodopsin C-tail from Cys322 to Ala348. On the left, colored bars indicate the portion of the C-tail visible in this structure (green), and in the arrestin-bound structure (salmon) (PDB id: 5W0P). On the right, the residue-residue contacts between rhodopsin C-tail and Gα (marked in green), Gβ (yellow), and arrestin (salmon) within 4 Å distance are indicated. Thr336 and Ser338 are phosphorylated in the arrestin-bound structure. The predicted phosphorylation sites are marked with red dots.

### Comparison of the GPCR-G protein binding interface

As in the other existing GPCR-G protein complex structures, the C-terminal α5 helix of Gα forms the major contact interface to rhodopsin. The α5 helix consists of the last 26 amino acids of Gα (H5.01-26), in which the last five residues fold into a hook-like structure (Tsai et al., 2018). The majority of the contacts formed by the α5 helix to GPCRs concentrate in the region from H5.11 to H5.26 (Glukhova et al., 2018) **(Suppl. Table 2)**.

We aligned the structures of available complexes using the Cα atoms of H5.11-26. This alignment reduces apparent differences in the binding poses and provides the “viewpoint of the G protein” **(Figure 3A)**. We compiled an exhaustive list of the residue-residue contacts between the receptor and the α5 helix in all the available GPCR/G protein complexes, and observed that the main contacts to the α5 helix are formed by TM5 and TM6, followed by TM3 and TM7/8 (**Figure 3B**, **Suppl. Table 3**).

**Figure 3.**
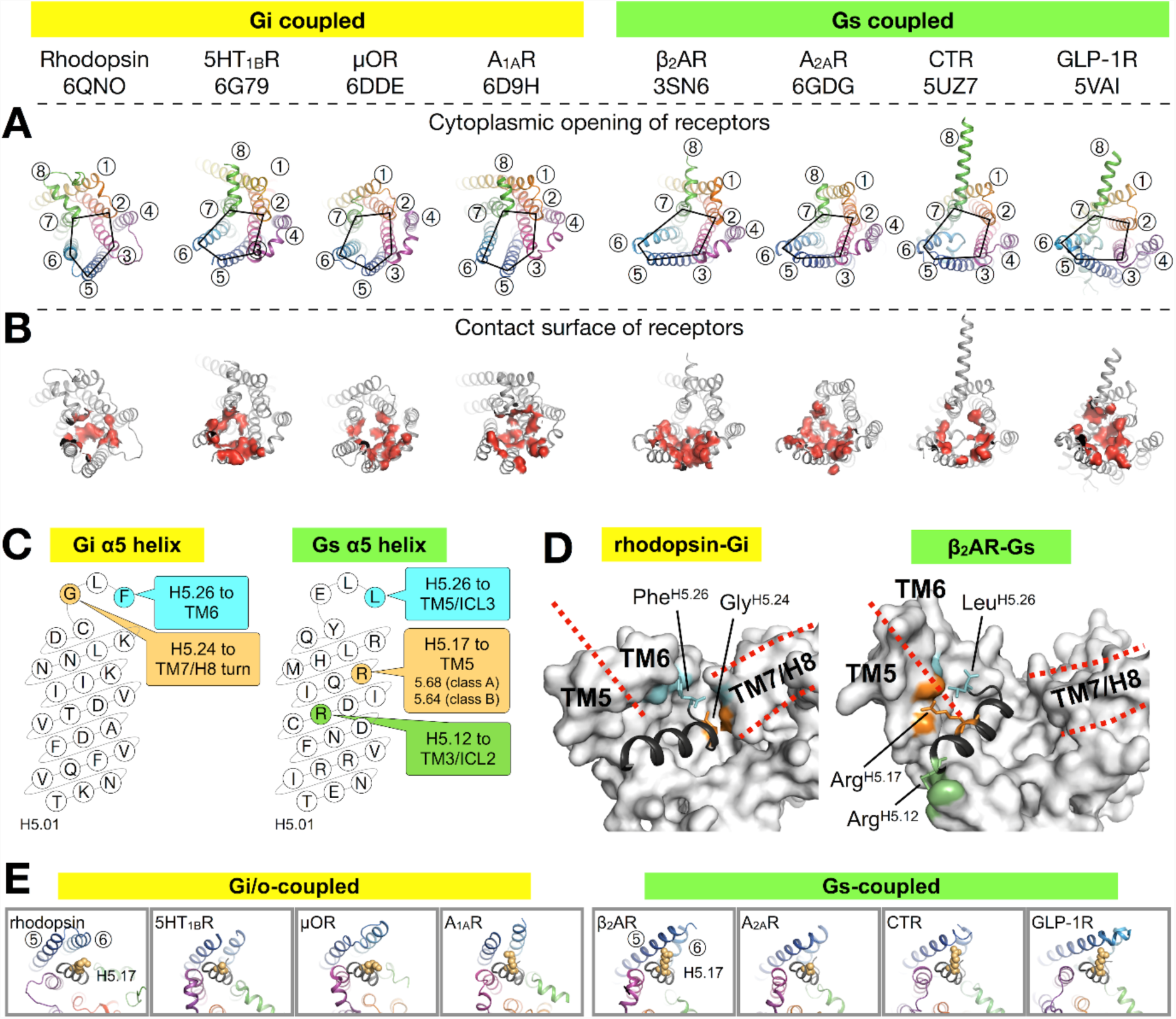
Binding of the Gα α5 helix in GPCR-G protein complexes. **(A)** Overview of a subset of GPCR-G protein complexes used for this analysis. For a complete list of the complexes used, see Suppl. Table 3. The complexes are shown from the cytoplasmic side, and the black pentagons connecting TM2-3-5-6-7 delineate the G protein-binding interface. Receptors are represented in ribbons with their TM helices numbered. **(B)** The red surfaces depict the contact area (within a distance of 4 Å) between the receptor and the α5 helix of Gα **(C)** Schematic representation of the α5 helix in Gi-and Gs-subtypes. The residues highlighted, and their respective binding site to the receptor, are conserved among all Gi, and Gs, complexes analyzed. All contacts retrieved from this analysis are reported in Suppl. Table 3. **(D)** Position of Gi-and Gs-specific contacts shown in the rhodopsin-Gi (this work, PDB id: 6QNO) and the β_2_AR-Gs (PDB id: 3SN6) complexes. The cytoplasmic side of the receptors is depicted as gray surface. Red lines mark the border between TM5 and TM6 and delineate H8. **(E)** Cytoplasmic view of GPCR-G protein complexes showing the interaction between H5.17 (orange spheres) and TM5 and TM6. From left to right, the structures (PDB ids) correspond to: this work, 6G79, 6DDE, 6D9H, 3SN6, 6GDG, 5UZ7, and 5VAI. Throughout the analysis, contacts are defined as atomic distances smaller than 4 Å.

Gi/o-bound complexes have two conserved contacts: one between Gly^H5.24^ and the TM7/H8 turn, and one between Tyr/Phe^H5.26^ and TM6. In Gs-bound complexes, three distinct interactions are found: between Arg^H5.12^ and TM3/ICL2, and between Leu^H5.26^ and Arg^H5.17^ and TM5 **(Figure 3C, D)**. In Gi/o complexes, the equivalent Lys^H5.17^ lies instead roughly parallel to TM6. Interestingly, Arg^H5.17^ appears to consistently interact with residue 5.68 in class A GPCRs, and with Lys^5.64^ in class B GPCRs **(Figure 3E, Suppl. Table 3)**. We suggest that a tight interaction between Arg^H5.17^ and TM5 might be one of the main determinants for the Gs-specific relocation of TM6.

Besides the canonical interaction with the α5 helix, our complex shows that ICL2 and ICL3 of the receptor are at contact distance to Gi (**Suppl. Figure 7A**). In all analyzed structures, we found that ICL2 lies near αN/β1 and β2/β3 of Gα, while ICL3 is close to α4/β6 **(Suppl. Figure 7 and Suppl. Figure 8)**. Interestingly, ICL2 folds into an α-helical structure in all the class A receptors except rhodopsin **(Suppl. Figure 9)**.

ICL2 does not contribute to binding Gα in the structures of 5HT_1B_-mini-Go and A_1A_R-G_i_. Nevertheless, the contact between Gα and ICL2 seems to discriminate between class A GPCRs – which interact via the αN/β1 loop– and class B GPCRs – which instead the region around β1 and β2/β3 **(Suppl. Fig. 8)**

ICL3, one of the most diverse regions in GPCRs, is often not completely resolved in the available structures (**Suppl. Fig. 7D**), either because it is too flexible or because it has been engineered to facilitate structural determination (Munk et al., 2019). Gi/o-coupled receptors use residues on TM5/ICL3/TM6 to contact the α4/β6 region, while Gs-coupled receptors mainly use TM5/ICL3 **(Suppl. Fig. 8)**. This difference is due to the larger displacement of TM6 in Gs complexes. In ICL3, residue Tyr^G.S6.02^, highly conserved in all G protein subtypes, engages the receptor in about half of all the complexes. This residue has been shown to be crucial for the stabilization of the rhodopsin-Gi complex (Sun et al., 2015).

Thus, our analysis suggests that ICL2 and ICL3 contribute to the binding interface between receptor and G protein, a feature that may be required to further stabilize the complex during nucleotide exchange.

## Discussion

In this work, we present the cryo-EM structure of the signaling complex between bovine rhodopsin and a Gi protein heterotrimer, stabilized using an antibody Fab fragment (Fab16) (Maeda et al., 2018). Overall, this structure agrees very well with existing complexes. In particular, we found that the binding mode of the G protein Ras domain – including the key C-terminal α5 helix– is virtually identical to our previously reported X-ray structure of the same receptor bound to a mini-Go protein (Tsai et al., 2018) **(Suppl. Fig. 6B)**.

However, our EM map shows a density on the Gβ subunit that extends from H8 of the receptor, constituting the proximal segment of the C-terminus (residues G324 to T335) **(Figure 2)**. One explanation of why the C-tail is observed in our density map and not in other structures may rely on the nature of the components used to reconstitute the reported GPCR complexes **(Suppl. Table 4)**. Among those, a meaningful comparison may be done with the μ-opioid receptor complex (Koehl et al., 2018), which is bound to a shorter version of our antibody Fab16, but in which the C-terminus of the receptor was partially truncated. In the human rhodopsin complex (Kang et al., 2018), which contains the full length C-tail, a recombinant Gβγ was used. Thus, our bovine rhodopsin complex contains the unique combination of a full-length C-tail, Gβγ purified from bovine rod outer segments, and Fab16 that may have contributed to trap the transient interaction of the intrinsically disordered C-tail to Gβ.

The Gβγ subunit engages with a wide range of effectors (Dupré, Robitaille, Rebois, & Hébert, 2009; Khan et al., 2013), including direct interactions with GPCRs. For instance, these can associate prior to trafficking to the plasma membrane (Dupré et al., 2006) where they can remain pre-coupled (Galés et al., 2005). Also, according to a sequential fit model (Herrmann et al., 2004, 2006), activated rhodopsin binds to Gβγ assisting to position the Ras domain of Gα into proximity of its binding site to the receptor. This would facilitate folding of the C-terminal α5 helix of this subunit leading to nucleotide exchange (Dror et al., 2015; Flock et al., 2015; Kapoor, Menon, Chauhan, Sachdev, & Sakmar, 2009). After dissociation of the G protein heterotrimer, the Gβ subunit recruits GRKs to the membrane, resulting in the phosphorylation of the receptor C-tail (Claing et al., 2002; Pao & Benovic, 2002; Pitcher et al., 1998) and binding of arrestin (Goodman Jr et al., 1996). Interestingly, HEK cells lacking functional Gα subunits (but retaining the native Gβγ) are still able to recruit arrestin, although they fail to activate ERK and whole-cell responses (Grundmann et al., 2018).

Our findings provide a structural explanation for these roles of Gβγ in GPCR-mediated signaling. We suggest that the observed interaction between the central part of the receptor C-terminus and Gβγ is involved in localizing the G protein heterotrimer to the active receptor first. After dissociation of the Gα subunit, a transient Gβγ/receptor complex could provide the adequate molecular context to allow receptor phosphorylation by bringing the kinase close to the receptor C-tail (Ribas et al., 2007).

In other class A GPCRs, interactions with Gβγ have been also located in ICL3 (in M2 and M3 muscarinic receptors, involved in the phosphorylation of this loop (Wu, Benovic, Hildebrandt, & Lanier, 1998) and potentially in ICL1 (in A_2_AR and β_2_AR) (García-Nafría, Lee, Bai, Carpenter, & Tate, 2018). Interestingly, the complexes of the class B GPCRs calcitonin and GLP-1 feature an extended and tilted H8 resulting in extensive contacts with Gβγ that have not been observed in Class A receptors (García-Nafría, Lee, et al., 2018). H8 has also been shown to bind Gβγ in the class B PTH1 receptor (Divieti, Mahon, Smrcka, & Bonacci, 2006). Thus, it is likely that class A and class B GPCRs use distinct mechanisms to engage Gβγ. The receptor-Gβγ interaction observed in our structure is compatible with proposed models of receptor-GRK recognition (Komolov et al., 2017; Sarnago et al., 2003). Thus, we suggest that the Gβγ subunit can ‘tag’ activated GPCRs and provide an interface for bringing different effectors (Gα subunits first, and GRKs later) near the cytoplasmic domains of GPCRs.

The availability of several GPCR-G protein complexes has greatly advanced our understanding on how receptors activate the Gα subunit (Dror et al., 2015; Flock et al., 2015). The binding interface of the receptor is partially formed by ICL2 and ICL3 (Chung et al., 2011; Glukhova et al., 2018; Sun et al., 2015). Accordingly, our EM map shows that these domains are in close proximity to Gi (**Suppl. Fig. 7A**). In particular, we found that ICL2 is at contact distance to the αN helix, the αN/β1 and the β2/β3 Gα in most structures (**Suppl. Fig. 7, Suppl. Fig. 8, and Suppl. Fig. 9**). While ICL2 contributes to the binding interface, there are no apparent conserved contacts among complexes (**Suppl. Fig. 8**). However, at the interface between receptor and the α5 helix of the G protein, in all Gi complexes Phe^H5.26^ and Gly^H5.24^ contact TM6 and TM7/H8 of the receptor, respectively. Instead, in all Gs complexes we find that Leu^H5.26^, Arg^H5.17^ and Arg^H5.12^ contact TM5/ICL3, TM5, and TM3/ICL2, respectively. In a recent analysis (Glukhova et al., 2018), it was proposed that residues 5.68 (class A GPCRs) and 5.64 (class B GPCRs) are particularly important for Gs protein binding. We observe that residue 5.68 interacts with H5.16 in both Gi-and Gs-bound complexes, but only with Arg^H5.17^ in all Gs complexes **(Suppl. Table 3)**. While our analysis is limited by the number of available structures, we suggest that the conserved contacts identified between α5 and the receptors are structurally important anchoring points, but we cannot exclude that they might be relevant at the level of recognition between G protein subtypes.

In order to visualize snapshots of the signaling pathway that have not been observed yet, more work on the structural characterization of complexes is still needed, ideally using the same receptor and different transducers. Such complexes are of pivotal importance to decipher the structural basis of the GPCR-mediated signaling cascade.

## Material and Methods

### Protein expression and purification

The N2C/M257Y/D282C mutant of bovine rhodopsin was expressed in human embryonic kidney (HEK) 293 GnTI^−^ cells as described (Deupi et al., 2012) (Deupi et al., 2012). The human Gαi subunit (Gαi1) with an N-terminal TEV protease-cleavable deca-histidine tag was expressed and purified as described (Sun et al., 2015). The transducin heterotrimer was isolated from the rod outer segment of bovine retina and Gβ1γ1 was separated from Gαt with Blue Sepharose 6 Fast Flow (GE Healthcare) as described (Maeda et al., 2014). The Gαi1β1γ1 heterotrimer (Gi) was prepared by mixing equimolar amounts of Gαi1 with or without 10xHis-tag and Gβ1γ1 and incubated at 4°C for 1 hour shortly before use for rhodopsin-Gi complex formation on the 1D4 immunoaffinity column.

### Fab16 production

The monoclonal mouse antibody IgG16 was generated as described (Koehl et al., 2018). Large scale production of IgG16 was performed using adherent hybridoma cell culture grown in DMEM medium supplemented with 10% ultra-low IgG fetal bovine serum (FBS) (Gibco, #16250078) and 25 U/ml of mouse interleukin-6 (Invitrogen, #PMC0064) at 37°C and 5% CO_2_. Antibody expression was increased by stepwise dilution of FBS concentration down to 2%. After incubation for 10-14 days, ∼500-ml cell suspension containing the secreted IgG was clarified by centrifugation and subsequent filtration through a 0.45 µm HAWP membrane (Merck Millipore). The filtrate was mixed with an equal volume of binding buffer (20 mM Na_2_HPO_4_/NaH_2_PO_4_, pH 7.0) and loaded to a 1-ml HiTrap Protein G Sepharose FF column (GE Healthcare). The column was washed with binding buffer until the UV_280_ absorbance dropped to a stable baseline, and IgG was eluted with 0.1 M glycine-HCl (pH 2.7). Fractions were immediately neutralized with 1 M Tris-HCl (pH 9.6). Fractions containing IgG16 were combined and dialyzed against 20 mM Na_2_HPO_4_/NaH_2_PO_4_ (pH 7.0), 1.5 mM NaN_3_ using a slide-A-lyzer dialysis cassette (12-14 kDa MWCO, Thermo Scientific) at 4 °C for 15 h. The dialysate was collected and mixed with the immobilized papain resin (0.05 ml resin for 1 mg IgG) (Thermo Scientific, #20341). Papain was activated by the addition of L-cysteine and EDTA to a final concentration of 20 mM and 10 mM respectively. IgG was digested overnight at 37°C with gentle mixing. Afterwards the immobilized papain resin was removed and the digested fraction was mixed with Protein A Sepharose (0.2 ml resin for 1 mg digested IgG, GE Healthcare, #17078001) for 1 hour at RT. Resins were washed with two column volumes (CV) of wash buffer 10 mM Tris (pH 7.5), 2.5 M NaCl. The flow-through and washing fractions containing Fab16 were collected and dialyzed against PBS supplemented with 1.5 mM NaN_3_ using a slide-A-lyzer dialysis cassette (12-14 kDa MWCO) at 4 °C. The dialysate was collected and concentrated with a VivaSpin 20 concentrator (10 kDa MWCO, Sartorius) to approximately 1.1 mg/ml. Glycerol was supplemented to the concentrated Fab16 at a final concentration of 10 %, and protein was flash frozen in liquid nitrogen and stored at −20 °C until use for formation of the rhodopsin-Gi-Fab16 complex.

### Fab16 crystallization and structure determination

For its crystallization, Fab16 was further purified by SEC on a Superdex 200 Increase 10/300 GL column (GE Healthcare) equilibrated in buffer 10 mM Tris (pH 7.5), 50 mM NaCl. Fractions containing pure Fab16 were collected and concentrated to approximately 14 mg/ml using a VivaSpin 4 concentrator (10 kDa MWCO, Sartorius). Fab16 was crystallized by vapor diffusion at 4 °C by dispensing 200 nl of protein and 200 nl of crystallization buffer containing 1.5 M malic acid (pH 7.5), 7 % (v/v) LDAO in an MRC 2-well crystallization plate (Swissci) using a mosquito^®^ crystallization robot. Crystals appeared after one day and grew to full size within 4 days. Crystals were soaked in the reservoir solution supplemented with 20 % (v/v) PEG 400 as a cryo-protectant and flash frozen in liquid nitrogen. The X-ray data was collected at the PXI beam line at the Swiss Light Source (SLS). Bragg peaks were integrated using XDS for individual datasets. XSCALE was used to scale and combine six datasets, and the pooled reflection list was further analyzed using the STARANISO server (Global Phasing Ltd.). The STARANISO server first analyzed the anisotropy for each dataset, giving a resolution of 1.90 Å in overall, 1.90 Å in the *0.92 a* - 0.38 c\** direction, 2.25 Å in the *k* direction, and 2.13 Å in the *-0.14 a* + 0.99 c\** direction under the criterion of CC½ = 0.3., following by scaling and merging of the reflections. The light chain from PDB id: 1MJU and the heavy chain from PDB id 2AAB without the CDR regions were used for molecular replacement using the program *Phaser-MR* in the Phenix suite. The model was built automatically using the Phenix *AutoBuild*, and the coordinates were manually adjusted using the visualization program Coot. The structure was refined using the *phenix.refine* to 1.90 Å **(Suppl. Table 4)**. The structure factor and the coordinates are deposited to the Protein Data Bank under the accession code 6QNK.

### Purification of the rhodopsin-Gi-Fab16 and the rhodopsin-G_i_ complexes

Buffers for every purification step were cooled to 4°C before use, and the steps after adding retinal were performed under dim red light before light activation of rhodopsin. The stabilized, constitutively active rhodopsin mutant N2C/M257Y/D282C was expressed in HEK293 GnTI^−^ deficient cells as described (Deupi et al., 2012). The cells were collected from the cell culture by centrifugation and homogenized in PBS buffer with cOmplete EDTA-free protease inhibitor (Roche). The cells were solubilized by supplementing dodecyl maltoside (DDM) (Anatrace, Sol-grade) to final concentrations of cell at 20 % (w/v) and of DDM at 1.25 % (w/v). After gentle stirring for 1-2 h, the solubilized fraction was collected after centrifugation at 200,000x g for 1 h. Solubilized rhodopsin apoprotein was captured in batch using the immunoaffinity Sepharose beads (GE Healthcare, #1043001) coupled to 1D4 antibody for more than 4 hour at the ratio of 5 g cells per ml resin. 1D4 resins were collected and washed with 10 CV of PBS, 0.04 % DDM. Afterwards, resins were resuspended in 2 CV of PBS, 0.04 % DDM and 75 µM of 9-cis (Sigma) or 11-cis retinal (Roche, or from NIH) for at least 6 hours in the dark. Later resins were washed with 30 CV of buffer A containing 20 mM HEPES pH 7.5, 150 mM NaCl, 1 mM MgCl_2_, 0.02 % (w/v) lauryl-maltose neopentyl glycol (LMNG) to wash off unbound retinal. Washed resins were resuspended in 1 CV of buffer A and mixed with 10xHis-tagged Gi heterotrimer at the ratio of 2 mg Gi per ml resin with the presence of 25 mU/ml apyrase (New England Biolabs). The resuspended resin/Gi mixture was subjected to irradiation with 495-nm long-passed filter for 10-15 minutes to induce activation of rhodopsin and G protein binding, followed by a 30-min incubation in the dark to allow full hydrolysis of nucleotide by apyrase. Resins were washed with 8 CV of buffer A to remove unbounded Gi heterotrimer. rhodopsin-G_i_ complex was eluted trice in batch with 1.5 CV of buffer containing 20 mM HEPES (pH 7.5), 150 mM NaCl, 1 mM MgCl_2_, 0.02 % LMNG, 80 µM 1D4 peptide (TETSQVAPA) for at least 2-hour incubation each time. Elution fractions were combined and incubated for at least 2-hour with Ni-NTA resins (0.5-2 ml), washed with 6 CV of 20 mM HEPES (pH 7.5), 150 mM NaCl, 50 mM imidazole, 0.01 % LMNG to remove free rhodopsin. Rhodopsin-Gi was eluted five times with 1 CV of 20 mM HEPES (pH 7.5), 150 mM NaCl, 350 mM imidazole, 0.01 % LMNG. Elution fractions were immediately concentrated using an Amicon Ultra concentrator (MWCO 100 kDa) with simultaneous buffer exchange to 20 mM HEPES (pH 7.5), 150 mM NaCl, 0.01 % LMNG. Rhodopsin-Gi was mixed with molar excess (1:1.4) of Fab16 and incubated for at least 1 h. The mixture of rhodopsin-Gi and Fab16 was concentrated using an Amicon Ultra concentrator (MWCO 30 kDa) and loaded to a Superdex 200 Increase 10/300 GL column for SEC with detergent-free buffer containing 20 mM HEPES (pH 7.5), 150 mM NaCl. Protein quality of each fraction was evaluated by UV-VIS measurement and SDS-PAGE. Fractions showing OD_280_/OD_380_ = ∼ 5.9 (ratio between 280 and 380 nm) was chosen for cryo-EM studies. For preparation of rhodopsin-Gi complex, purified rhodopsin and 10xHis-tag-free Gi heterotrimer were mixed in a test tube in equimolar ratio with the presence of 25 mU/ml of apyrase and incubated for 1 hour at 4°C. The protein mixture was concentrated and purified using a Superdex 200 Increase 10/300 GL column. Fractions showing OD_280_/OD_380_ = 3 were used to prepare cryo-EM specimens.

### Cryo-electron microscopy and image analysis

Purified samples of rhodopsin-Gi with and without Fab16 were plunge-frozen in a Vitrobot MarkIV (FEI Company) operated at 4°C and 90% humidity. A drop of 3.5 µL sample at 0.2 mg/mL was dispensed onto a glow discharged lacey carbon grid (Ted Pella, Inc.) and blotted for 2-3 seconds before vitrification in liquid ethane. Images were acquired by a Titan Krios microscope operated at 300 kV equipped with a Falcon III, or with a K2 Summit and GIF energy filter **(Suppl. Table 1)**. Datasets were pre-processed and pruned in FOCUS (Biyani et al., 2017) using MotionCor2 (Zheng et al., 2017) with dose weighting for movie alignment, and CTFFIND4 (Rohou & Grigorieff, 2015) for micrograph contrast transfer function estimation. Automated particle picking was performed in Gautomatch (http://www.mrc-lmb.cam.ac.uk/kzhang/Gautomatch/) and all further processing steps were carried in cryoSPARC (Structura Biotechnology Inc.) and RELION 2 and 3 (Kimanius, Forsberg, Scheres, & Lindahl, 2016; Zivanov et al., 2018). Around 115,000 particles from the three best resolved 3D classes were pooled and subjected to 3D auto-refinement with a soft mask which deliberately excludes the density of the AH domain observed in one of the 3D classes. Map sharpening and modulation transfer function (MTF) correction were performed with RELION post-processing. The resulting map has a nominal resolution of 4.38 Å estimated following the gold standard Fourier Shell Correlation (FSC) at FSC = 0.143. Local resolution estimation was performed using *blocres* (Cardone, Heymann, & Steven, 2013).

### Model building and structure refinement

The initial models of rhodopsin and the Ras-like domain of Gαi protein were adapted from the structure of rhodopsin-mini-Go complex (PDB id: 6FUF). The initial model of Gβγ was obtained from the crystal structure of guanosine 5’-diphosphate-bound transducin (PDB id: 1GOT). The models were docked into the 3D map as rigid bodies in Chimera (Pettersen et al., 2004). The coordinates of the structure were manually adjusted and the C-terminus of rhodopsin was built in Coot (Emsley & Cowtan, 2004), which was used to visualize the EM map and the models. The models were refined using the *phenix.real_space_refine* in the Phenix suite (Adams et al., 2010).

### Comparison and analysis of structures

Structural models were downloaded from the Protein Data Bank. For comparing structures of the GPCR-G protein complexes, the residue-residue contacts within 4 Å were first identified using PyMOL (The PyMOL Molecular Graphics System, Version 2.0 Schrödinger, LLC).

### Figure preparation

Figures were prepared using ChimeraX (Goddard et al., 2018) and PyMOL (The PyMOL Molecular Graphics System, Version 2.0 Schrödinger, LLC).

## Acknowledgments

We thank ScopeM at ETH Zurich for support, and in particular to Mr. Peter Titmann for help with the operation of Titan Krios for the Falcon 3 datasets. We are indebted to Dr. Mohamed Chami from the Center for Cellular Imaging and NanoAnalytics in Basel for his help with the initial screening of cryo-grids. We thank Dr. Takashi Ishikawa and the Electron Microscopy Facility of PSI for his support. We thank Jean-Philippe Carralot (F. Hoffmann-La Roche Ltd) for help in antibody generation, Martin Siegrist, Georg Schmid, Bernard Rutten, Doris Zulauf, Stephanie Kueng (Roche Non-Clinical Biorepository) and Ralf Thoma for technical assistance for biomass and cell line generation.

## Author contributions

G.F.X. Schertler designed and supervised the project. S. Maeda and C.J. Tsai characterized Fab16 and its use for the production of the rhodopsin complex. F. Pamula purified the complexes for cryo-EM studies. J. Mühle, F. Pamula, and C.J. Tsai crystallized and solved the structure of Fab16. R. Dawson coordinated large-scale cell culture, and developed Fab16, with help from H. Matile. J. Marino coordinated the cryo-EM part of the project, acquired EM data with help of ScopeM and C-CINA, and contributed to EM analysis. R.J. Adaixo performed most EM data processing, with help from H. Stahlberg. I. Mohamed helped in setting up acquisition of a Falcon3 dataset at ETH Zurich. C.J. Tsai, T. Flock, N. Taylor and X. Deupi modeled the complex into the EM map. C.J. Tsai performed the structural analysis on GPCR complexes. J. Marino, C.J. Tsai, and X. Deupi analyzed the data and wrote the manuscript, with help from all co-authors.

## Data availability

The cryo-EM density map was deposited under accession code EMD-4598 on the EM Data Bank. The related structure coordinates of the rhodopsin-complex bound to Fab16 (accession code 6QNO) and the crystal structure of Fab16 (accession code 6QNK) were deposited on the Protein Data Bank.

