## Supplementary Material for "Cryo-EM structure of the rhodopsin-Gαi-βγ complex reveals binding of the rhodopsin C-terminal tail to the Gβ subunit"

**Supplementary Figure 1. Purification of the rhodopsin-Gi and rhodopsin-Gi-Fab16 complexes. (A)** SDS-PAGE analysis of individual proteins and main peak fraction of corresponding size-exclusion chromatography (SEC). **(B)** SEC profiles of rhodopsin-Gαi and rhodopsin-$G_i$-Fab16 complexes used for the preparation of cryo-grids.

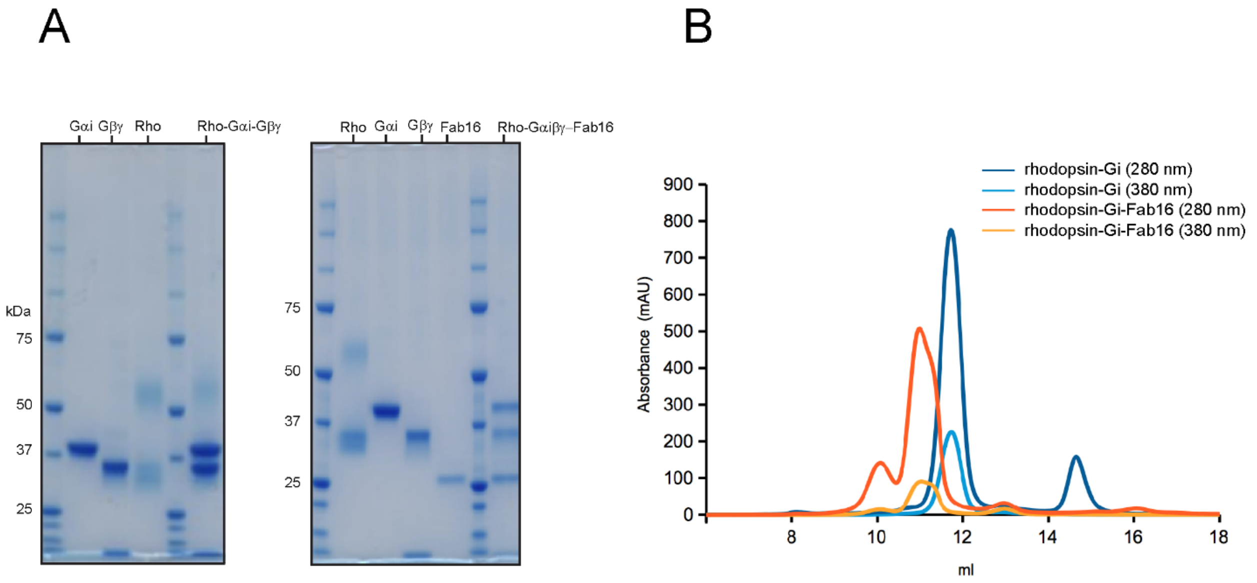

**Supplementary Figure 2. Cryo-EM maps of rhodopsin-$G_i$ complexes with and without Fab16. (A)** Small dataset of rhodopsin-Gi (without Fab16) obtained using a Falcon III detector. **(B)** Small dataset of rhodopsin-Gi-Fab16 obtained using a Falcon III detector. **(C)** Gold-standard FSC curves with resolutions estimated at the 0.143 cut-off for the density maps of the two datasets.

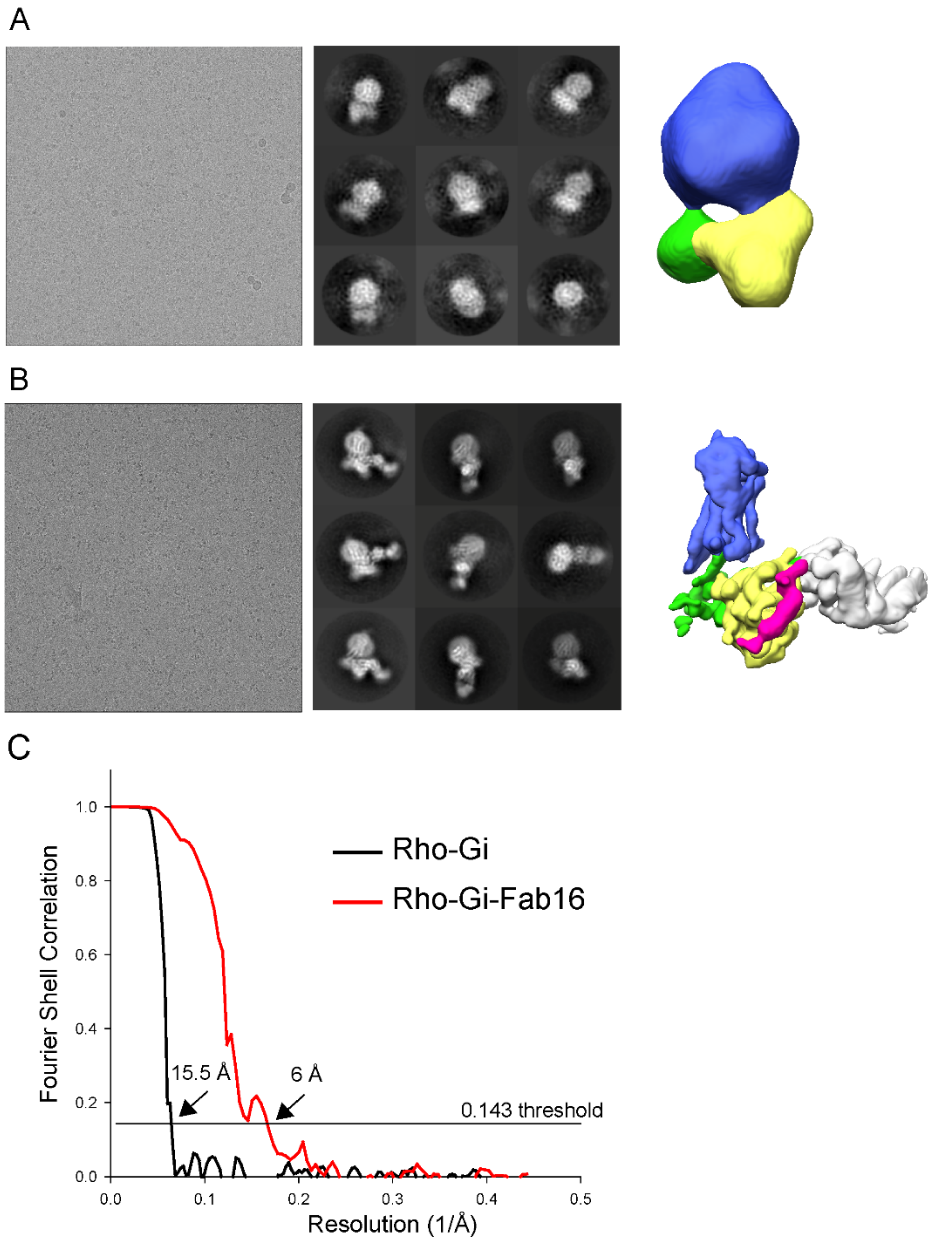

**Supplementary Figure 3. Image processing of the rhodopsin-Gi-Fab16 complex acquired with K2. (A)** A representative image. **(B)** 2D class averages. **(C)** Image-processing workflow of the data processing. **(D)** Local resolution of the 3D reconstruction. **(E)** Fourier Shell Correlation plot of the two half datasets generated in RELION.

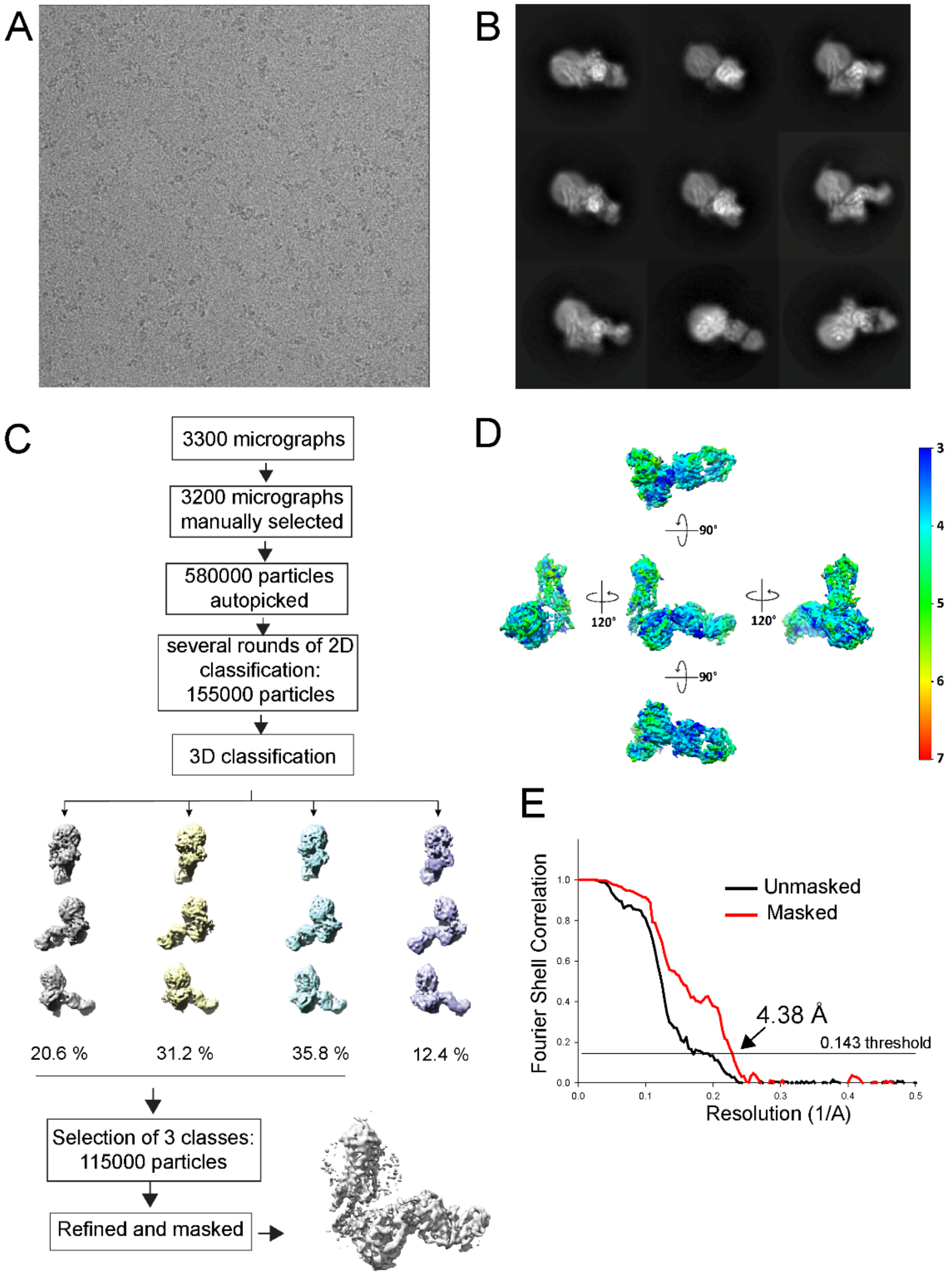

**Supplementary Figure 4. 3D classification reveals the flexibility of the AH domain of Gαi. (A)** Density map of one 3D class obtained during classification in RELION. The AH domain of the Gα subunit, highlighted in red, becomes visible only at high threshold when visualizing the density map in Chimera. **(B)** Overview of the three density maps obtained during 3D classification. The region corresponding to the AH domain of Gαi, which displays higher heterogeneity in the three density maps, is indicated with a red circle.

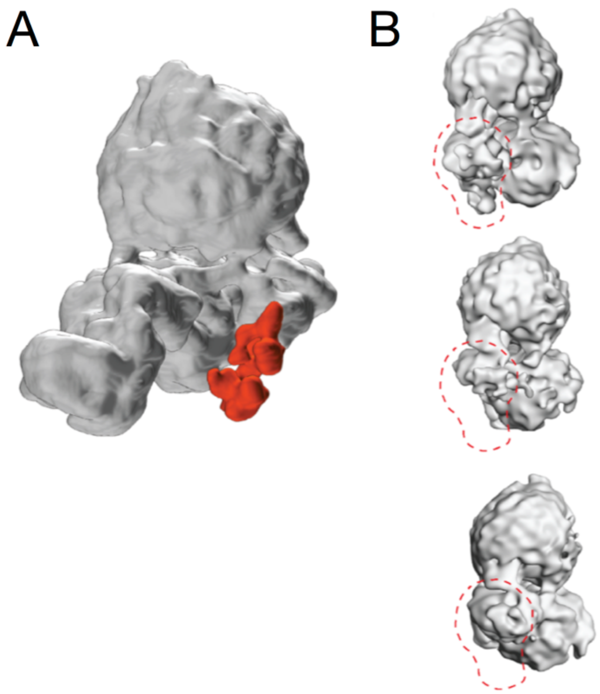

**Supplementary Figure 5. Details of the cryo-EM density map of rhodopsin-$G_i$-Fab16 with a fitted atomic model. (A)** The segmented density map shows individual regions of rhodopsin (transmembrane helices (TM), intracellular and extracellular loops (ICL, ECL), helix 8 (H8) and the C-terminal tail (C-tail), retinal-binding pocket; rhodopsin – blue, retinal – orange) and Gαi (αN and α5 helices, Gαi in green). Atoms are colored by type (nitrogen – blue; oxygen – red; sulfur – dark yellow). **(B)** Binding interface between the α5 helix of Gαi (in green) and the TM helices of rhodopsin (in blue). The maps are plotted at a 10 σ cuf-off using Pymol.

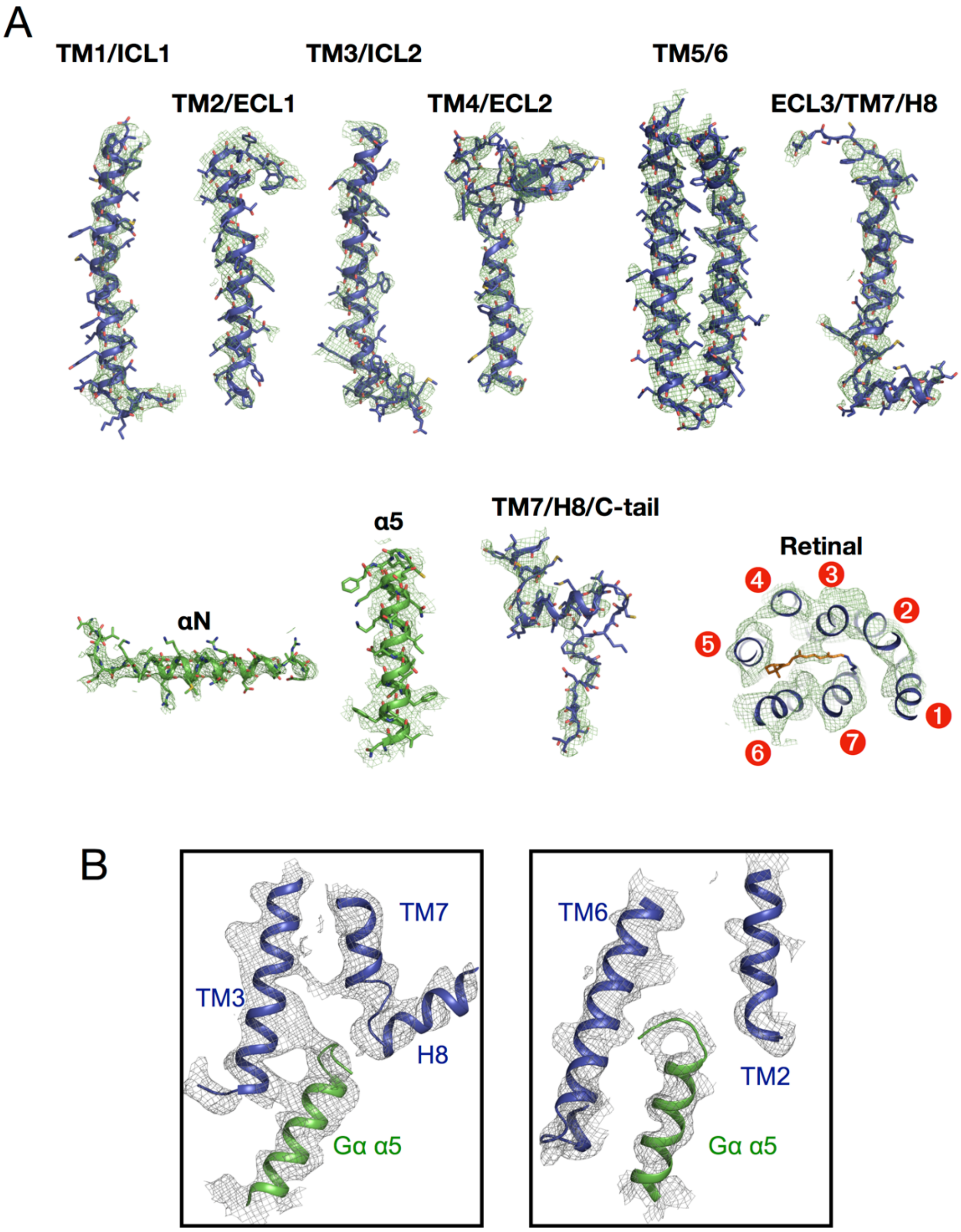

**Supplementary Figure 6. Comparison of the bovine rhodopsin-Gi complex to the other GPCR-G protein complexes. (A)** The complex structures are aligned to the Cα atoms of the rhodopsin-Gi complex (receptor – blue; Gα – green; Gβ – yellow; Gγ – magenta; peptide ligand – salmon). **(B)** Binding interface between the receptor and the Gα α5 helix in the structures of rhodopsin-Gi and rhodopsin-mini-Go (PDB id: 6FUF). Rhodopsin and G proteins are colored in blue and green, respectively. The side chains of Leu131$^{3.46}$, Arg135$^{3.50}$, Tyr306$^{7.53}$ are plotted in sticks and the density maps are shown as grey meshes.

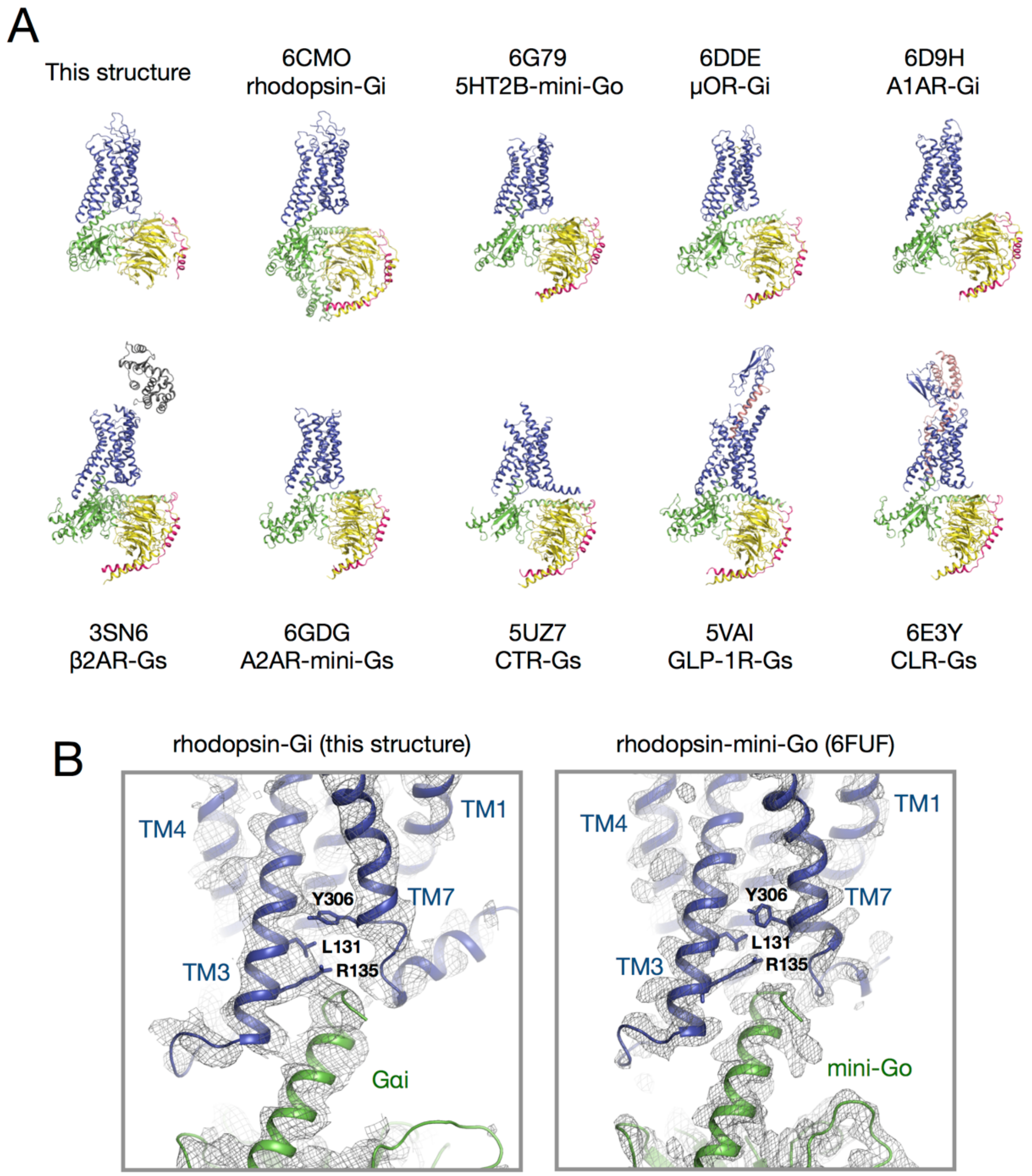

**Supplementary Figure 7. Contacts observed between ICL2/ICL3 and the Gα subunit. (A)** Intracellular loop (ICL) 2 and 3 in the EM density map **(B)** Schematic overview of the contacts between the intracellular loops of the receptor and Gα. **(C)** Sequence of ICL2 in the available structures of the GPCR-G protein complexes. Residues that contact to regions of the G protein other than the α5 helix are highlighted in cyan. **(D)** Same as panel B, but for ICL3. Residues missing from the structure are marked in grey. The residues are numbered using the Ballesteros-Weinstein scheme.

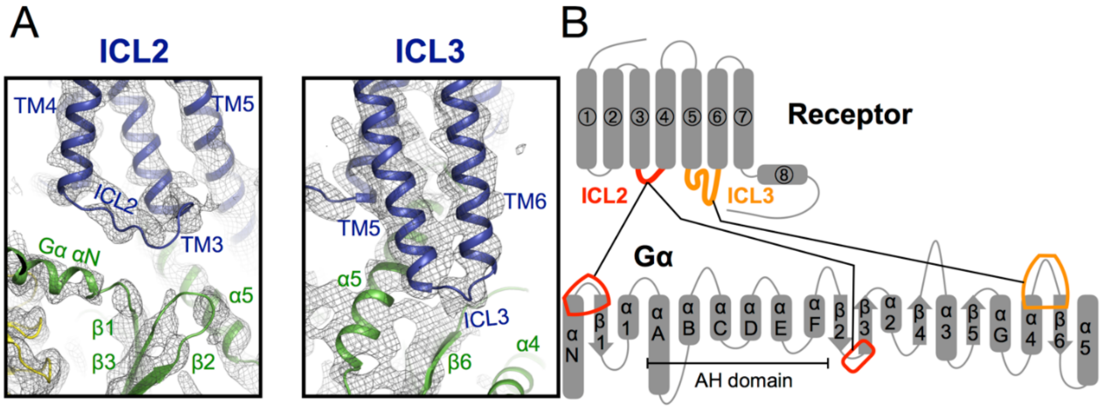


C

| Class | Coupling | | Receptor | 3.55 | 3.56 | 3.57 | 3.58 | 3.59 | 3.60 | 34.50 | 34.51 | 34.52 | 34.53 | 34.54 | 34.55 | 34.56 | 34.57 | 4.37 | 4.38 | 4.39 | 4.40 | 4.41 |
|---|---|---|---|---|---|---|---|---|---|---|---|---|---|---|---|---|---|---|---|---|---|---|
| | | | | TM3 | | | | | | ICL2 | | | | | | | | TM4 | | | | |
| Class A | $G_{i/o}$-coupled | This structure | bovine Rho | C | K | – | – | – | – | P | M | S | N | F | R | F | – | – | G | E | N | H |
| Class A | $G_{i/o}$-coupled | 6FUF | bovine Rho | C | K | – | – | – | – | P | M | S | N | F | R | F | – | – | G | E | N | H |
| Class A | $G_{i/o}$-coupled | 6CMO | human Rho | C | K | – | – | – | – | P | M | S | N | F | R | F | – | – | G | E | N | H |
| Class A | $G_{i/o}$-coupled | 6DDE or 6DDF | μOR | C | H | – | – | – | – | P | V | K | A | L | D | F | R | – | T | P | R | N |
| Class A | $G_{i/o}$-coupled | 6G79 | 5HT1BR | T | D | – | – | – | – | A | V | E | Y | S | A | K | R | – | T | P | K | R |
| Class A | $G_{i/o}$-coupled | 6D9H | A1AR | K | I | – | – | – | – | P | L | R | Y | K | M | V | V | – | T | P | R | R |
| Class A | $G_{i/o}$-coupled | 6N4B | CB1R | H | R | – | – | – | – | P | L | A | Y | K | R | I | V | – | T | R | P | K |
| Class A | $G_s$-coupled | 3SN6 | β2AR | T | S | – | – | – | – | P | F | K | Y | Q | S | L | L | – | T | K | N | K |
| Class A | $G_s$-coupled | 5G53 | A2AR | R | I | – | – | – | – | P | L | R | Y | N | G | L | V | – | T | G | T | R |
| Class A | $G_s$-coupled | 6GDG | A2AR | R | I | – | – | – | – | P | L | R | Y | N | G | L | V | – | T | G | T | R |
| | | | | | | | | | | ICL2 tentative numbering | | | | | | | | | | | | |
| Class B | $G_s$-coupled | 5UZ7 | CTR | H | T | L | I | V | V | A | V | F | T | E | K | Q | – | – | – | – | – | R |
| Class B | $G_s$-coupled | 5VAI | GLP-1R | Y | T | L | L | A | F | A | V | F | – | – | – | – | – | S | E | Q | R | I |
| Class B | $G_s$-coupled | 6B3J | GLP-1R | Y | T | L | L | A | F | S | V | L | S | – | – | – | – | – | E | Q | W | I |
| Class B | $G_s$-coupled | 6E3Y | CLR | H | T | L | I | V | V | A | V | F | A | E | K | Q | – | – | – | – | – | H |

D

| Class | Coupling | | Receptor | 5.65 | 5.66 | 5.67 | 5.68 | 5.69 | 5.70 | 5.71 | 5.72 | 5.73 | 5.74 | 5.75 | 5.76 | 56.01 | 56.02 | 56.03 | 56.04 | 56.05 | 56.06 | 56.07 | 56.08 | 6.21 | 6.22 | 6.23 | 6.24 | 6.25 | 6.26 | 6.27 | 6.28 | 6.29 | |
|---|---|---|---|---|---|---|---|---|---|---|---|---|---|---|---|---|---|---|---|---|---|---|---|---|---|---|---|---|---|---|---|---|---|
| | | | | TM5 | | | | | | | | | | | | ICL3 | | | | | | | | TM6 | | | | | | | | | |
| | | | | | | | | | | | | | | | | ICL3 tentative numbering | | | | | | | | | | | | | | | | | |
| Class A | $G_{i/o}$-coupled | This structure | bovine Rho | V | K | E | A | A | A | Q | Q | – | – | – | – | Q | E | S | – | – | – | – | – | – | – | – | A | T | T | Q | K | A | |
| Class A | $G_{i/o}$-coupled | 6FUF | bovine Rho | V | K | E | A | A | A | Q | Q | – | – | – | – | Q | E | S | – | – | – | – | – | – | – | – | A | T | T | Q | K | A | |
| Class A | $G_{i/o}$-coupled | 6CMO | human Rho | V | K | E | A | A | A | Q | Q | – | – | – | – | Q | E | S | – | – | – | – | – | – | – | – | A | T | T | Q | K | A | |
| Class A | $G_{i/o}$-coupled | 6DDE or 6DDF | μOR | L | K | S | V | – | – | – | – | – | – | – | – | R | M | L | S | G | – | – | – | – | – | S | K | E | K | D | R | N | |
| Class A | $G_{i/o}$-coupled | 6G79 | 5HT1BR | A | R | S | R | I | L | K | Q | T | P | – | – | N245 – K302 | | | | | | | | – | – | – | K | L | M | A | A | R | |
| Class A | $G_{i/o}$-coupled | 6D9H | A1AR | I | R | K | Q | L | N | K | K | V | – | – | – | S216 – G220 | | | | | | | | – | D | P | Q | K | Y | Y | G | K | |
| Class A | $G_{i/o}$-coupled | 6N4B | CB1R | | | | | | | | | | | R | G | T | Q314-R331 | | | | | | | – | – | – | P | D | Q | A | R | M | |
| Class A | $G_s$-coupled | 3SN6 | β2AR | A | K | R | Q | L | Q | K | I | D | K | S | E | G | R | F240 – S261 | | | | | | – | – | – | S | K | F | C | L | K | |
| Class A | $G_s$-coupled | 5G53 | A2AR | A | R | R | Q | L | K | Q | M | E | S | | | Q214 – G218 | | | | | | | | E | R | A | R | S | T | L | Q | K | |
| Class A | $G_s$-coupled | 6GDG | A2AR | A | R | R | Q | L | K | Q | M | E | S | – | – | Q214 – G218 | | | | | | | | E | R | A | R | S | T | L | Q | K | |
| Class B | $G_s$-coupled | 5UZ7 | CTR | M | R | – | – | – | – | – | – | – | – | – | – | E | T | H | E332 – H336 | | | | M | – | – | – | – | – | – | – | – | – | Y6.35 |
| Class B | $G_s$-coupled | 5VAI | GLP-1R | L | K | A | – | – | – | – | – | – | – | – | – | N | L | M | C | K | T | D | I | – | – | – | – | – | – | – | – | – | K6.35 |
| Class B | $G_s$-coupled | 6B3J | GLP-1R | L | K | A | – | – | – | – | – | – | – | – | – | N338 – T343 | | | | | | | | – | – | – | – | – | – | – | – | – | D6.33 |
| Class B | $G_s$-coupled | 6E3Y | CLR | L | K | – | – | – | – | – | – | – | – | – | – | V | T | H324 – S328 | | | | N | L | – | – | – | – | – | – | – | – | – | Y6.35 |

**Supplementary Figure 8. Residue-residue contacts between ICL2/3 and Gα.** Contacts at the regions of ICL2 and 3 are identified using a 4-Å cut-off. Residue-residue contacts are listed for the regions of ICL2 (top panel) and ICL3 (bottom panel). The residues of Gα are numbered following the common Gα protein numbering (CGN). Contacts observed in the individual structures are noted as colored bullet symbols (blue – Gi-coupled receptors; green – Gs-coupled, class A receptors; magenta – Gs-coupled, class B receptors).

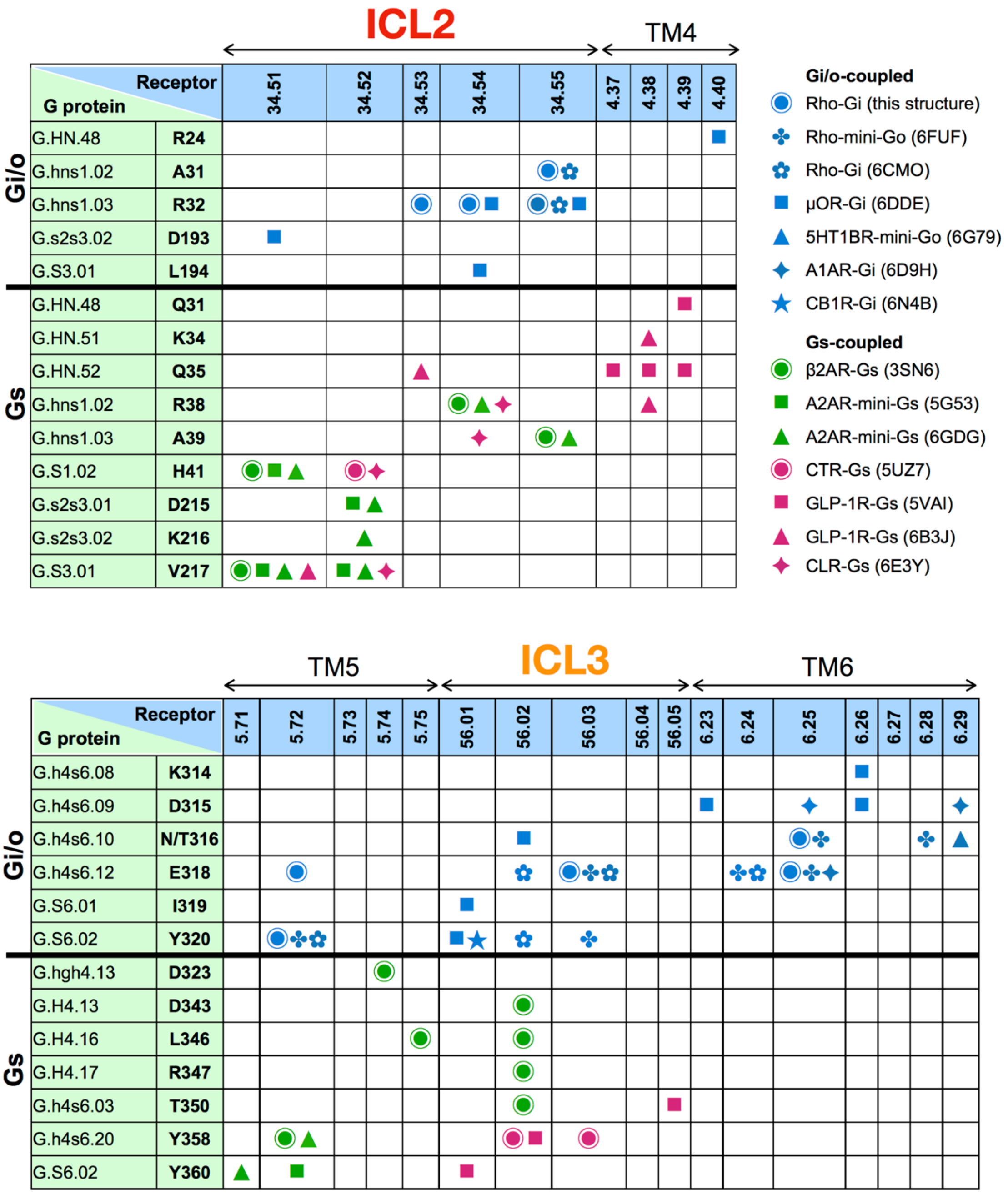

**Supplementary Figure 9. Region near ICL2 in available structures.** View of the existing GPCR-G protein complex structures centered at the ICL2 region. The structures are aligned to the Cα atoms of rhodopsin. The regions of the G protein involved in contacting ICL2 are the αN/β1 and β2/β3 turns, and the α5 helix.

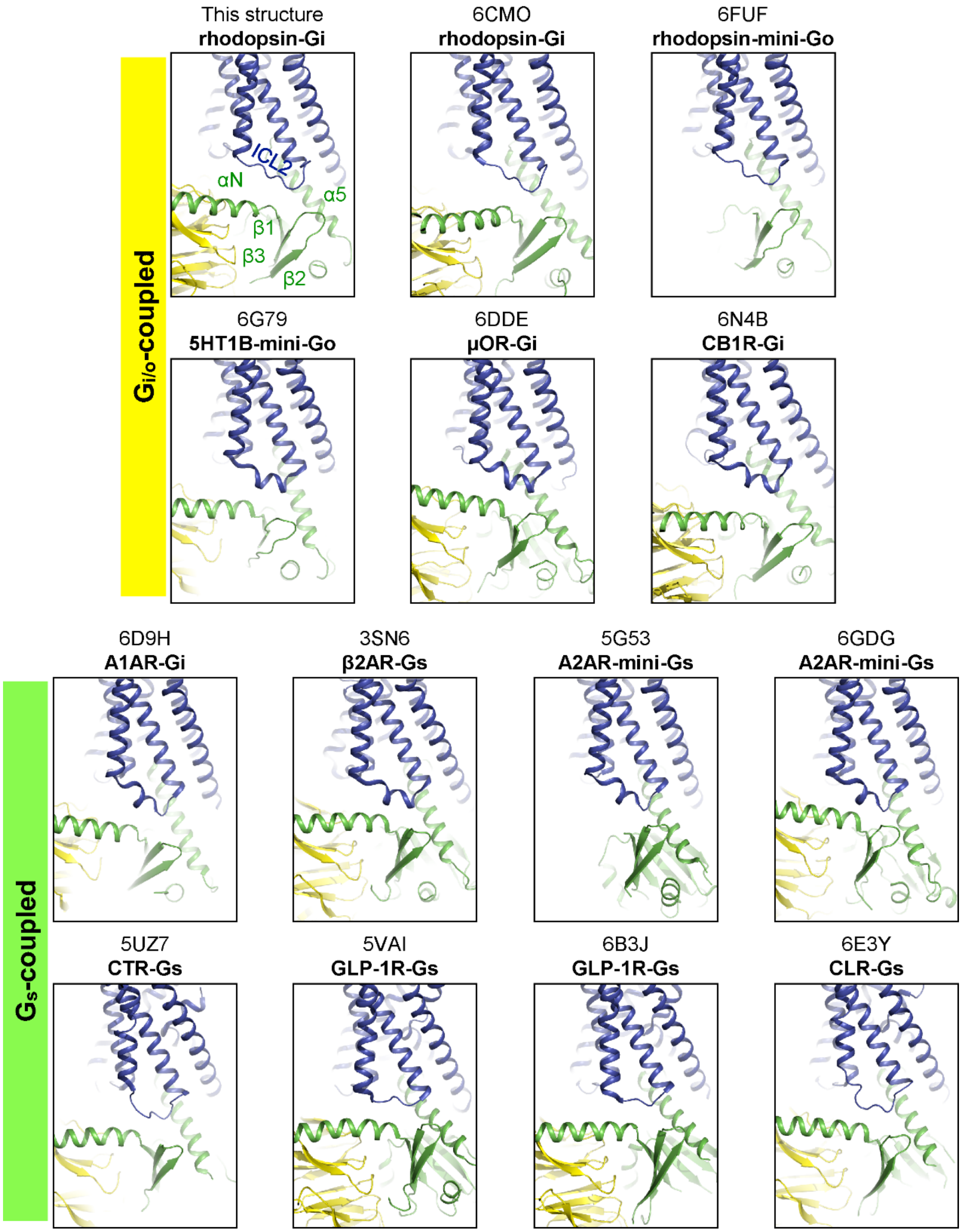

**Supplementary Table 1. Cryo-EM data collection and refinement statistics.**

| | **Dataset #1 (-) Fab** | **Dataset #2 (+) Fab** | **Dataset #3 (+) Fab** |
|---|---|---|---|
| Direct Electron Detector | Falcon III (EC mode) | Falcon III (EC mode) | K2 Summit + GIF (super resolution mode) |
| N. of movies after cleaning | 600 | 600 | 3200 |
| Accelerating Voltage (kV) | 300 | 300 | 300 |
| Total Exposure (e/Å$^2$) | 50 | 50 | 60 |
| Number of frames per movie | 50 | 50 | 40 |
| Defocus Range (µm) | -1.5 to -2.5 | -1.5 to -2.5 | -1.5 to -2.5 |
| Automated EM data acquisition software | EPU | EPU | SerialEM |
| Pixel Size (Å/px) | 1.12 | 1.12 | 0.83 |
| Symmetry | C1 | C1 | C1 |
| N. of particles in final map | 69000 | 65000 | 115000 |
| Final Map Resolution (Å) | 15.5 | 6 | 4.38 |
| Map Sharpening B-factor (Å$^2$) | -- | -380 | -175 |

**Supplementary Table 2. Residue-residue contact list between GPCRs and Gα H5.11-26 within 4 Å.**

| Gi/o coupled | Receptor | H5.01 | H5.02 | H5.03 | H5.04 | H5.05 | H5.06 | H5.07 | H5.08 | H5.09 | H5.10 | H5.11 | H5.12 | H5.13 | H5.14 | H5.15 | H5.16 | H5.17 | H5.18 | H5.19 | H5.20 | H5.21 | H5.22 | H5.23 | H5.24 | H5.25 | H5.26 | Number of atomic contacts |
|---|---|---|---|---|---|---|---|---|---|---|---|---|---|---|---|---|---|---|---|---|---|---|---|---|---|---|---|---|
| | | Residues of Gα α5 helix | | | | | | | | | | | | | | | | | | | | | | | | | | |
| This structure | bovine Rho | T | K | N | V | Q | F | V | F | D | A | V | T | D | V | I | I | K | N | N | L | K | D | C | G | L | F | |
| 6CMO | human Rho | T | K | N | V | Q | F | V | F | D | A | V | T | D | V | I | I | K | N | N | L | K | D | C | G | L | F | |
| 6FUF (mini-Go) | bovine Rho | T | N | N | A | Q | V | I | F | D | A | V | T | D | I | I | I | A | N | N | L | R | G | C | G | L | Y | |
| 6DDE | μOR | T | K | N | V | Q | F | V | F | D | A | V | T | D | V | I | I | K | N | N | L | K | D | C | G | L | F | 1 |
| 6DDF | μOR | T | K | N | V | Q | F | V | F | D | A | V | T | D | V | I | I | K | N | N | L | K | D | C | G | L | F | 2-4 |
| 6G79 (mini-Go) | 5HT1BR | T | N | N | A | Q | V | I | F | D | A | V | T | D | I | I | I | A | N | N | L | R | G | C | G | L | Y | 5-7 |
| 6D9H | A1AR | T | K | N | V | Q | F | V | F | D | A | V | T | D | V | I | I | K | N | N | L | K | D | C | G | L | F | 8-10 |
| 6N4B | CB1R | T | K | N | V | Q | F | V | F | D | A | V | T | D | V | I | I | K | N | N | L | K | D | C | G | L | F | 11-13 |

| Gs coupled | Receptor | H5.01 | H5.02 | H5.03 | H5.04 | H5.05 | H5.06 | H5.07 | H5.08 | H5.09 | H5.10 | H5.11 | H5.12 | H5.13 | H5.14 | H5.15 | H5.16 | H5.17 | H5.18 | H5.19 | H5.20 | H5.21 | H5.22 | H5.23 | H5.24 | H5.25 | H5.26 | Number of atomic contacts |
|---|---|---|---|---|---|---|---|---|---|---|---|---|---|---|---|---|---|---|---|---|---|---|---|---|---|---|---|---|
| | | Residues of Gα α5 helix | | | | | | | | | | | | | | | | | | | | | | | | | | |
| 3SN6 | β2AR | T | E | N | I | R | R | V | F | N | D | C | R | D | I | I | Q | R | M | H | L | R | Q | Y | E | L | L | |
| 5G53 (mini-Gs) | A2AR | T | E | N | A | R | R | I | F | N | D | C | R | D | I | I | Q | R | M | H | L | R | Q | Y | E | L | L | |
| 6GDG (mini-Gs) | A2AR | T | E | N | A | R | R | I | F | N | D | C | R | D | I | I | Q | R | M | H | L | R | Q | Y | E | L | L | 1 |
| 5UZ7 | CTR | C | E | N | I | R | R | V | F | N | D | C | R | D | I | I | Q | R | M | H | L | R | Q | Y | E | L | L | 2-4 |
| 5VAI | GLP-1R | T | E | N | I | R | R | V | F | N | D | C | R | D | I | I | Q | R | M | H | L | R | Q | Y | E | L | L | 5-7 |
| 6B3J | GLP-1R | F | T | N | I | R | R | V | F | N | D | C | R | D | I | I | Q | R | M | H | L | R | Q | Y | E | L | L | 8-10 |
| 6E3Y | CLR | – | – | N | I | R | R | V | F | N | D | C | R | D | I | I | Q | R | M | H | L | R | Q | Y | E | L | L | 11-13 |

Number of atomic contacts

sequence of α5 helix

### Supplementary Table 3. Residue-residue contacts between GPCRs and Gα H5.11-26.

| Gα α5: | H5.11 | H5.12 | H5.13 | H5.15 | H5.16 | H5.17 | H5.19 | H5.20 | H5.22 | H5.23 | H5.24 | H5.25 | H5.26 |
|---|---|---|---|---|---|---|---|---|---|---|---|---|---|
| **Gi/o coupled** | | | | | | | | | | | | | |
| Rho: This structure | - | - | Q237-5.72 | - | - | - | V138-3.53 | V139-3.54 | L72-2.39 | L72-2.39<br>R135-3.50<br>V138-3.53 | R135-3.50<br>N310-8.47 | R135-3.50<br>M257-6.40 | T242-6.25<br>K245-6.28<br>A246-6.29<br>E249-6.32<br>K311-8.48 |
| Rho: 6CMO | - | - | - | - | - | S240-ICL3 | - | - | - | - | M309-7.56<br>N310-8.47<br>K311-8.48 | R135-3.50<br>E249-6.32<br>M253-6.36 | T242-6.25<br>K245-6.28<br>A246-6.29<br>K311-8.48 |
| Rho: 6FUF | - | - | Q237-5.72<br>S240-ICL3<br>T243-6.26 | - | A233-5.68 | - | V138-3.53 | V139-3.54<br>A246-6.29<br>V250-6.33 | L72-2.39 | R135-3.50<br>N310-8.47 | N310-8.47<br>K311-8.48 | R135-3.50<br>M257-6.40 | T242-6.25<br>K245-6.28<br>A246-6.29<br>K311-8.48 |
| 5HT1BR: 6G79 | - | V155-ICL2 | R238-5.68 | - | I151-3.43<br>A154-ICL2<br>R238-5.68 | - | A150-3.53<br>I151-3.54 | I151-3.54<br>A235-5.65<br>R238-5.68 | S372-7.56<br>N373-8.47 | R147-3.50<br>S372-7.56 | K311-6.32<br>T315-6.36<br>S372-7.56 | I231-5.61<br>A312-6.33 | R238-5.68<br>I239-5.69<br>R308-6.29<br>K311-6.32 |
| µOR: 6DDE | - | P172-ICL2 | V262-5.68<br>R263-ICL3<br>M264-ICL3 | P172-ICL2<br>L176-ICL2 | V169-3.54<br>P172-ICL2<br>R258-5.64 | M264-ICL3 | A168-3.53<br>R179-ICL2 | V169-3.54<br>L259-5.65 | T103-2.39 | T103-2.39<br>R165-3.50 | D340-8.47 | R277-6.32<br>M281-6.36 | I278-6.33 |
| A1AR: 6D9H | - | - | Q210-5.68<br>K213-5.71 | P112-ICL2<br>L113-ICL2 | P112-ICL2<br>Q210-5.68 | L211-5.69 | R108-3.53<br>P112-ICL2 | V109-3.43<br>I207-5.65 | D42-2.37<br>F45-2.40<br>R108-3.53<br>I292-8.47<br>K294-8.49 | R105-3.50<br>R108-3.53<br>I292-8.47 | R291-7.56 | R105-3.50<br>V203-5.61<br>I207-5.65<br>I232-6.33<br>L236-6.37 | K228-6.29 |
| CB1R: 6N4B | - | - | - | L222-ICL2 | P221-ICL2 | - | S217-3.53<br>P221-ICL2<br>Y224-ICL2 | S217-3.53<br>I218-3.54 | R150-2.37<br>S152-2.39 | S152-2.39 | S401-8.47 | R214-3.50<br>L341-6.33 | M337-6.29<br>R340-6.32 |
| **Gs coupled** | | | | | | | | | | | | | |
| β2AR: 3SN6 | F139-ICL2 | T136-3.55<br>P138-ICL2<br>F139-ICL2<br>K140-ICL2 | Q229-5.68<br>K232-5.71 | P138-ICL2<br>Q142-ICL2 | I135-3.54<br>P138-ICL2<br>E225-5.64<br>R228-5.67<br>Q229-5.68 | Q229-5.68<br>I233-5.72 | A134-3.53<br>Y141-ICL2<br>Q142-ICL2 | I135-3.54<br>A226-5.65<br>Q229-5.68 | - | R131-3.50 | K270-6.32<br>T274-6.36 | I135-3.54<br>V222-5.61<br>A226-5.65<br>A271-6.33<br>T274-6.36<br>L275-6.37 | L230-5.69 |
| A2AR: 5G53 | L110-ICL2 | R107-3.55<br>I108-3.56<br>P109-ICL2<br>L110-ICL2 | Q207-5.68<br>Q210-5.71 | P109-ICL2 | I106-3.54<br>P109-ICL2<br>A203-5.64<br>Q207-5.68 | Q207-5.68 | A105-3.53<br>I106-3.54<br>Y112-ICL2 | I106-3.54<br>A204-5.65 | I292-8.47 | R102-3.50<br>A105-3.53<br>R291-7.56 | R291-7.56<br>R293-8.48<br>R296-8.51 | I200-5.61<br>A231-6.33<br>L235-6.37<br>R291-7.56 | L208-5.69<br>K227-6.29 |
| A2AR: 6GDG | L110-ICL2 | R107-3.55<br>P109-ICL2<br>L110-ICL2 | Q207-5.68 | P109-ICL2 | I106-3.54<br>P109-ICL2<br>Q207-5.68 | Q207-5.68 | A105-3.53<br>I106-3.54<br>Y112-ICL2 | I106-3.54<br>A204-5.65 | I292-8.47<br>R293-8.48 | R102-3.50 | S234-6.36<br>R291-7.56<br>R293-8.48 | I200-5.61<br>A231-6.33<br>S234-6.36<br>L235-6.37 | L208-5.69<br>K227-6.29 |
| CTR: 5UZ7 | - | V252-ICL2<br>K326-5.64 | - | V252-ICL2 | I248-3.58<br>V249-3.59<br>V252-ICL2<br>K326-5.64 | K326-5.64 | L247-3.57<br>T254-ICL2 | L323-5.61 | R180-2.46 | R180-2.46<br>Y243-3.53<br>L244-3.54 | L348-6.45<br>C394-7.60 | T345-6.42<br>L348-6.45 | M327-5.65 |
| GLP-1R: 5VAI | - | A256-3.59 | K334-5.64 | - | L255-3.58 | K334-5.64 | L254-3.57 | V331-5.61 | R176-2.46<br>E408-8.49 | H180-2.50<br>L251-3.54<br>L356-6.45<br>L359-6.48 | S352-6.41<br>L401-7.56<br>Y402-7.57<br>V405-7.60<br>N406-8.47<br>N407-8.48 | S352-6.41<br>T353-6.42 | L339-ICL3 |
| GLP-1R: 6B3J | - | - | K334-5.64 | S258-ICL2 | L255-3.58<br>K334-5.64 | K334-5.64 | L254-3.57<br>L255-3.58 | L255-3.58<br>V331-5.61 | R176-2.46 | Y250-3.53<br>L251-3.54 | N407-8.48 | S352-6.41<br>L356-6.45 | - |
| CLR: 6E3Y | - | V242-3.59<br>V243-3.60<br>V245-ICL2<br>F246-ICL2 | K319-5.64 | V245-ICL2<br>F246-ICL2 | I241-3.58<br>V242-3.59<br>V245-ICL2<br>K319-5.64 | K319-5.64 | L240-3.57 | I241-3.58<br>L316-5.61 | R173-2.46 | R173-2.46<br>Y236-3.53<br>L237-3.54 | R336-6.40<br>F387-7.60<br>N388-8.47<br>G389-8.48 | L316-5.61<br>R336-6.40<br>A337-6.41<br>L341-6.45 | L316-5.61<br>L320-5.65<br>K333-6.37<br>R336-6.40 |

**Supplementary Table 4. Details of the source organism of the Gα, Gβ, and Gγ proteins used to obtain GPCR G-protein complexes for structure determination.**

| Complex | PDB | Gα | Uniprot ID | Gβ | Uniprot ID | Gγ | Uniprot ID |
|---|---|---|---|---|---|---|---|
| **bovine Rho-Gi** | this work | Human Gαi1 | P63096 | Bovine Gβ1 | P62871 | Bovine Gγ1 | P02698 |
| **human Rho-Gi** | 6CMO | Human Gαi1 | P63096 | Rat Gβ1 | P54311 | Bovine Gγ2 | P63212 |
| **μOR-Gi** | 6DDE | Human Gαi1 | P63096 | Human Gβ1 | P62873 | Human Gγ2 | P59768 |
| **5HT1BR-mini-Go** | 6G79 | Human Gαo1 (mini) | P09471 | Human Gβ1 | P62873 | Human Gγ2 | P59768 |
| **A1AR-Gi** | 6D9H | Human Gαi2 | P04899 | Human Gβ1 | P62873 | Human Gγ2 | P59768 |
| **CB1R-Gi** | 6N4B | Human Gαi1 | P63096 | Human Gβ1 | P62873 | Human Gγ2 | P59768 |
| **β2AR-Gs** | 3SN6 | Bovine Gαs1 | P04896 | Rat Gβ1 | P54311 | Bovine Gγ2 | P63212 |
| **A2AR-mini-Gs** | 6GDG | Human Gαs2 (mini) | P63092 | Human Gβ1 | P62873 | Human Gγ2 | P59768 |
| **CTR-Gs** | 5UZ7 | Human Gαs2 | P63092 | Human Gβ1 | P62873 | Human Gγ2 | P59768 |
| **GLP-1R-Gs** | 5VAI | Human Gαs2 | P63092 | Rat Gβ1 | P54311 | Bovine Gγ2 | P63212 |
| **GLP-1R-Gs** | 6B3J | Human Gαs2 | P63092 | Human Gβ1 | P62873 | Human Gγ2 | P59768 |
| **CLR-Gs** | 6E3Y | Human Gαs2 | P63092 | Human Gβ1 | P62873 | Human Gγ2 | P59768 |

**Supplementary Table 5. Crystallographic data and structural refinement of Fab16**

| **Data collection** | |
|---|---|
| Space Group | C 1 2 1 |
| Cell dimensions *a, b, c* (Å) | 169.94, 69.08, 133.76 |
| *α, β, γ* (°) | 90.0, 127.2, 90.0 |
| Wavelength (Å) | 1.0 |
| Resolution (Å) | 47.54 – 1.90 (2.05 – 1.90) |
| $R_{pim}$ | 0.019 (0.457) |
| $R_{merge}$ | 0.086 (2.121) |
| $I/\sigma I$ | 19.8 (1.4) |
| Completeness (%) | 95.1 (86.0) |
| Multiplicity | 20.8 (22.3) |
| CC1/2 | 1.000 (0.701) |

| **Refinement statistics** | |
|---|---|
| Refinement program | PHENIX 1.13_2998 |
| Resolution | 47.56 – 1.90 (1.968 – 1.9000) |
| No. Reflections | 67762 (1294) |
| $R_{work}/R_{free}$ (%) | 17.95 (36.11)/21.22 (36.11) |
| Number of atoms | |
| Total | 7313 |
| Protein | 6771 |
| Tetraethylene glycol (PG4) | 39 |
| 1,2-ethanediol (EDO) | 48 |
| Triethylene glycol (PGE) | 20 |
| Malic ion (MLT) | 36 |
| Water | 390 |
| Ramachandran favoured (%) | 98.27 |
| Ramachandran allowed (%) | 1.73 |
| Ramachandran outliers (%) | 0.00 |
| R.m.s.d Bond legth (Å) | 0.005 |
| R.m.s.d Bond angles (°) | 1.14 |
| Averaged B factor ($Å^2$) | |
| Protein | 50.13 |
| Non-water solvent | 80.09 |
| Water | 52.23 |
